# Plant selection and ecological microhabitat drive domestications of shrub-associated microbiomes in a revegetated shrub ecosystem

**DOI:** 10.1101/2023.01.19.524707

**Authors:** Zongrui Lai, Yanfei Sun, Yang Yu, Zhen Liu, Yuxuan Bai, Yangui Qiao, Lin Miao, Weiwei She, Shugao Qin, Wei Feng

**Affiliations:** Yanchi Research Station, School of Soil and Water Conservation, Beijing Forestry University, Beijing 100083, China; Key Laboratory of Genetics and Germplasm Innovation of Tropical Special Forest Trees and Ornamental Plants, Ministry of Education, College of Forestry, Hainan University, Haikou, 570228, China; CAS Engineering Laboratory for Yellow River Delta Modern Agriculture, Institute of Geographic Sciences and Natural Resources Research, Chinese Academy of Sciences, Beijing, 100101, China

**Keywords:** dryland ecosystem, plant microbiomes, community assembly, plant-microbe interactions, co-occurrence networks

## Abstract

Shrubs are used for revegetation of degraded dryland ecosystem worldwide and could recruit large numbers of microbes from the soil; however, the plant-associated microbiome assembly and the effect of plant introduction on the soil microbiomes are not fully understood. We detected shrub-associated microbes from five ecological microhabitats, including the leaves, litter, roots, rhizosphere, and root zone, across four xeric shrub plantations (*Artemisia ordosica, Caragana korshinskii, Hedysarum mongolicum*, and *Salix psammophila*). To detect the patterns of shrub-associated microbiome assembly, 16S and ITS2 rRNA gene sequencing was performed. PERMANOVA and differential abundance analysis demonstrated that changes in the bacterial and fungal communities were more dependent on the microhabitats rather than on the plant species, with distinct niche differentiation. Moreover, source tracking and nestedness analysis showed that shrub-associated bacteria were primarily derived from bulk soils and slightly pruned in different microhabitats; however, a similar pattern was not found for fungi. Furthermore, the surrounding zone of roots was a hotpot for microbial recruitments of revegetated shrubs. Null model analysis indicated that homogeneous selection of determinism dominated the bacterial communities, whereas dispersal limitation and undominated process of stochasticity drove the assembly of fungal communities. Our findings indicate that ecological microhabitat of revegetated shrublands was the main predictor of the bacterial and fungal compositional variances. This study will help advance our understanding of the mechanism underlying the plant-soil microbiome feedbacks during the initial plant-establishment period in a dryland ecosystem. Further, this work provides theoretical reference for establishment and sustainable management of shrublands in drylands.

## 1. Introduction

Shrubs are foundation species in fragile dryland ecosystems and performs diverse ecological functions, such as maintaining species diversity, controlling soil erosion, and promoting soil formation (Maestre et al., 2021). Each plant taxon harbours a characteristic mixture of microorganisms, collectively termed as the ‘plant microbiota’ (Xu et al., 2021; Vandenkoornhuyse et al., 2015). These plant-specific microbiomes impact the nutrition and water acquisition of xeric plants, suppress diseases, and support plant health and resistance to harsh environments (i.e., drought, salt, and high temperature) (Cordovez et al., 2019; Trivedi et al., 2020); thereby, driving plant-soil feedbacks (Delgado-Baquerizo et al., 2020). Drylands, covering approximately 40% of the Earth’s land surface, are expanding globally due to climate warming (Berg and McColl, 2021) and contain ubiquitous microbial species (Soussi et al., 2016). Many studies have focused on certain traits of soil microbes in drylands, such as the diversity, phyletic classification, and biogeography (Maestre et al., 2015; Soussi et al., 2016; Sun et al., 2020); however, the interplay between desert plants and soil microbiomes remains largely elusive, which limits the understanding of plant-soil feedback. Further, an improved understanding of the plant-soil microbiomes can improve degraded land restoration programs and increase the productivity of desert ecosystems in the future (Trivedi et al., 2019).

Although the soil microbiome is a common reservoir of microbes, plants (i.e., leaves, stem, and roots) provide diverse microhabitats for colonization by numerous microorganisms (Beckers et al., 2017). Plant-associated microbiomes acting as a second genome has received substantial attention in the recent years (Berg et al., 2014; Turner et al., 2013). Several studies have demonstrated that plant-associated microbial composition and functions are regulated by plant host genotype (Beckers et al., 2017; Soussi et al., 2016; Vandenkoornhuyse et al., 2015), i.e., each plant habitat harbours its own characteristic microbiome (Cordovez et al., 2019). Moreover, host plants can provide a number of habitats for microorganisms and then subtly affect microbiomes that confer tolerance to abiotic stress by supplying photosynthetic carbon (Müller et al., 2016; Xu et al., 2021). For example, root exudate and rhizodeposition predominantly influence the rhizosphere microbiota (Gupta et al., 2021; Zhalnina et al., 2018). In the phyllosphere, cytokinins drive the assembly and function of microbiomes (Hu et al., 2018). Recent studies indicate that plant microbiome assembly and function are profoundly affected by different seasons or plant developmental stages (Xiong et al., 2021a; Aleklett et al., 2022). In drylands, plants thrive under prolonged environmental stresses, such as high irradiance, drought, and salt accumulation, through the development of specific physiological and molecular extremophile traits (VanWallendael et al., 2019; Van Zelm et al., 2020). Previous studies have shown that large plant microbiomes, special root-associated bacteria, and fungi are also beneficial for desert plants in coping with unfavourable conditions (Soussi et al., 2016; Liu et al., 2021). Nevertheless, the mechanisms underlying the development of dryland plant microbiome remain very limited, impeding our understanding of desert ecological functions and processes.

Recently, niche differentiation of plant-associated microbial taxa, mainly at the soil-root interface (rhizosphere and root endosphere), has received great research attention (Trivedi et al., 2019; Xiong et al., 2021a; Wang et al., 2022). In fact, each plant tissue (fruits, seeds, flowers, leaves, stems, and roots) and soil habitat (rhizosphere and litter) provide unique ecological niche that supports a characteristic microbial community (Gupta et al., 2021; Vandenkoornhuyse et al., 2015; Xu et al., 2021). Potentially, different microhabitats reflect different biotic (substrate and organic matter) and abiotic (temperature and water availability) conditions (Müller et al., 2016; Zheng and Gong., 2019). Compared to other humid ecosystems, dryland ecosystems have greater differences in biotic and abiotic conditions across microhabitats (Soussi et al., 2016; Trivedi et al., 2019). However, such interactions in dryland ecosystems remain to be elucidated.

Furthermore, recent studies suggest that the formation and development of microbial communities across microhabitats are not only controlled by environmental factors but are also strongly interrelated with each other (Zheng and Gong, 2019; Bernard et al., 2021; Walsh et al., 2021). Generally, members of the microbiome are horizontally acquired from the surrounding environments where the initial and main reservoir is the soil (Cordovez et al., 2019; Xiong et al., 2021a), while others migrate vertically via parents of the host plants (Vandenkoornhuyse et al., 2015). The belowground plant compartments harbour more microbes than aboveground plant tissues (Zheng and Gong, 2019), and the rhizosphere, a 1 mm thin zone of soil that surrounds fine roots, is more enriched with microbes than the bulk soil (Philippot et al., 2013). Thus, plant-associated microbes are selectively recruited by the plants, and the plant compartment defines the composition of the microbiomes. Taxonomic and genomic analyses have shown that there are overlapping microbial communities among different plant compartments (Cordovez et al., 2019; Edwards et al., 2015). For example, although the leaves and roots of *Arabidopsis thaliana* (L.) Heynh. have specific microbiota members, they have a part of similar functional diversities (Bai et al., 2015). Whether microbial functional overlap is attributed to the migration of microorganisms among the different compartments, and whether these compartment microbiome assemblies are mainly influenced by the soil microbiome, remain to be elucidated (Turner et al., 2013; Xu et al., 2021). Therefore, clarifying the diversity, abundance, composition, and dynamics of each microhabitat is helpful for improving the understanding of plant-environment interactions (Bulgarelli et al., 2012).

In drylands, woody plants are widely revegetated and shrubs, being the frontier species, are frequently adopted worldwide to combat desertification and to maintain sand dune stability (Zastrow, 2019). Notably, revegetation not only improve the microenvironment, but also increase plant material input to soil (Arneth et al., 2021); however, the effect of revegetation on the assembly of plant-associated and soil microbiome should be explored. In 2011, permanent plots (*Artemisia ordosica, Caragana korshinskii, Hedysarum mongolicum*, and *Salix psammophila* plantations) were established in the study site. We found that the effect of fine roots on soil organic carbon varied across the shrublands, and soil microbial diversity and composition were also significantly different (Lai et al., 2016; Liu et al., 2018; Sun et al., 2020). Therefore, a multi-plant experiment was performed in the same permanent plots. An adjacent bare sandy land, land before revegetation, served as a control in this study. We aimed to observe: (1) a drastic microbial community differentiation among the four revegetated shrubs; (2) distinct bacterial and fungal communities and compositions across different plant species and microhabitats (phylloplane, detritusphere, root rhizoplane, rhizosphere soil, and root zone soil) and the largest microbial species reservoir found in bulk soil. Consequently, we tried to identify the mechanisms involved in the assembly processes of the plant-associated microbiomes.

## 2. Materials and methods

### 2.1. Field experiment and sampling

In 2001, four desert shrub populations, which include *A. ordosica, C. korshinskii, H. mongolicum*, and *S. psammophila*, were planted on bare sandy land at the Yanchi Research Station (37° 42’ 31’’ N/107° 13’ 47’’ E, 1,530 m above sea level, shrubland details see Lai et al. 2016), located in the Mu Us Desert, Ningxia, China were used in this study. The study area was fenced to avoiding livestock grazing and anthropogenic disturbance. The long-term mean annual temperature was 8.1 °C, ranging from -8.4- 22.7 °C, and the mean annual precipitation was 292 mm at this study site (Liu et al., 2018). All selected shrubs were synchronously planted in the same field, which was characterised by sandy soil and subjected to the same management practices (Sun et al., 2020). The field experiments (four shrubland plots and one bare sandy land) were performed in August 2018 (Table S1), when the shrubs grow vigorously. For sampling, twelve 10 m × 10 m plots were randomly selected in each shrublands. The distance between plots was range from 10 m to 100 m. The leaves (matured, 15 g), detritus (twig and leaf litter, 15 g), fine roots (< 2 mm, 15 g), soil from the rhizosphere (surrounding the fine roots, 50 g) and the root zone (under the canopy, 50 g) from three healthy shrubs in each plot (samples from three shrubs were mixed and formed a pooled sample), and bulk soil (50 g) from bare sandy land were carefully collected in a single day, using disposable gloves to avoid contamination. Root and soil samples were collected from four soil profiles (40 cm × 40 cm × 40 cm) around the shrub at a distance of 0.2 - 1.0 m. Bulk soils from the same depth were randomly collected from the bare sandy land plot adjacent to other plots. Twelve samples per sample type (soil and plant) were collected for two days (10–11 August). All samples were stored separately in sealed 50-mL centrifuge tube, immediately transported to the Magigene Biotechnology Lab (Guangzhou, Guangdong Province) on dry ice within 48 h and store at -70 °C until further molecular analyses.

### 2.2. DNA extraction and sequencing

All frozen samples were transported on dry ice to maintain a temperature below 4 °C for DNA extraction and sequencing as soon as possible after field sampling (within a week). Visible soil debris in plant tissues (leaf, detritus, and root) were washed using distilled water. Then, approximately 1 g of crushed plant tissue and 0.5 g of soil sample were used to extract DNA using the MoBio PowerSoil^®^ DNA Isolation Kit (MoBio Laboratories, Inc., Carlsbad, CA, USA) and DNA samples were placed randomly across plates. The concentration and purity of all extracts were measured using the NanoDrop One (Thermo Fisher Scientific, MA, USA) and quantified again prior to polymerase chain reaction (PCR) (Agler et al., 2016).

A two-step barcoded PCR protocol was used to maximise the phylogenetic coverage of bacteria and fungi (Bai et al., 2015; Lundberg et al., 2013). Primers for the tagging the bacterial and fungal amplicons were 515F/806R (515F: 5′-GTGCCAGCMGCCGCGGTAA; 806R: 5′-GGACTACHVGGGTWTCTAAT) and ITS5-1737F/ITS2-2043R (ITS5-1737F: GGAAGTAAAAGTCGTAACAAGG; ITS2-2043R: GCTGCGTTCTTCATCGATGC), respectively, and were used in equal concentrations (Ihrmark et al., 2012; Kembel et al., 2014). After PCR amplification, the length and concentration of amplicons were detected using 1% agarose gel electrophoresis. The PCR products were purified using the EZNA^®^ Gel Extraction Kit (Omega Bio-Tek, Doraville, USA). Sequencing libraries were generated using NEBNext^®^ Ultra™ DNA Library Prep Kit for Illumina^®^ (New England Biolabs, MA, USA). Illumina MiSeq sequencing was carried out on the IlluminaHiseq2500 platform (Illumina Inc., San Diego, CA, USA) using 2×250 bp.

Quality filtering of paired-end raw reads and assembly of paired-end clean reads were performed using Trimmomatic v0.33 (http://www.usadellab.org/cms/?page=trimmomatic) and FLASH v1.2.11 (https://ccb.jhu.edu/software/FLASH/), respectively (Durán et al., 2018). Raw tag quality control was analysed using Mothur V1.35.1 (http://www.mothur.org) (Schloss et al., 2009). Operational taxonomic unit (OTU) clustering, species annotation, and phylogenetic relationship construction were performed using a combination of USEARCH v10 (Caporaso et al., 2010), KRONA, GraPhlAn, QIIME v1.9.1, and R v3.6.3 software. Contaminant sequences (e.g., protista, archaea, chloroplast, mitochondrial, and viridiplantae sequences) were filtered from the data set (Edgar, 2010). Sequences were assigned to taxonomy based on a 97% sequence similarity threshold (Gunnigle et al., 2017). In total, 17, 954, 236 bacterial and 14, 207, 200 fungal high-quality raw reads from 252 samples. A total of 10,000 and 20,000 reproducible and measurable OTUs for bacteria and fungi, respectively, were included in the complete datasets, and the full dataset was split into phylloplane, detritusphere, root rhizoplane, rhizosphere, root zone soil, and bulk soil sub-datasets to examine the differences among the shrub species (DeSantis et al., 2006). Samples from the microhabitats (i.e., leaf, detritus, root, and soil) were rarefied separately to minimise sample loss (bacteria: 34528 reads; fungi: 28374 reads). All analyses conducted had six replicates. All DNA-sequencing data were uploaded to the NCBI Sequence Read Archive (SRA) with the accession number SRP348383.

### 2.3. Statistical analyses

All the data and Figureures were run in the R statistical software v3.6.3 (The R Foundation for Statistical Computing, Vienna, Austria; http://www.r-project.org). Data normality was examined using the Shapiro–Wilk rank sum test. PROC UNIVARIATE was used to test normality of distribution and homogeneity of variance for residuals. To satisfy the homoscedasticity assumption, OTUs were normalised using variance-stabilizing transformation. Statistical significance was determined at α = 0.05, and when necessary, *P* values for multiple comparisons were corrected using sequential Bonferroni correction. The differential abundance, richness, and α-diversity across species and microhabitats were identified using two-way ANOVA models (the aov R function). For each ANOVA model, multiple comparisons were FDR-corrected. Significant differences between shrub species or microhabitats were evaluated with the Kruskal-Willis rank sum test (kruskal.test with dunn.test in R; FDR-corrected *p* < 0.05). Normality of the diversity data was checked with the Shapiro-Wilk test. If the data was skewed, log_10_-transformed data were used to statistical analysis. Differential abundances for bacteria and fungi in each shrubland and microhabitat compared with bare sandy land were determined using DESeq2 (Love et al., 2014), with FDR-corrected *p* < 0.05 considered significant.

To determine the differences in the microbial community, Bray-Curtis dissimilarity matrices were calculated and then visualised with non-metric dimensional scaling (NMDS) ordinations. Permutational multivariate analysis of variance (PERMANOVA) pairwise comparisons were conducted using the adonis function in the R package vegan with 999 permutations for bacteria and fungi for statistically supporting the visual clustering results of the NMDS analyses (Oksanen et al., 2019). The co-occurrence network was constructed using the IGRAPH package in R, based on Spearman’s rank correlations of all OTUs, accompanied by the calculations of the descriptive and topological network properties (Hartman et al., 2018), and visualised the significant correlations (Spearman’s *r* > 0.6 or *r* < -0.6, *p* < 0.01) in GEPHI v.0.9.2 (https://gephi.org/). The average degree (the number of direct correlations to a node) is defined as the network complexity.

SourceTracker, based on Bayesian approach, was performed to evaluate the source of the plant microbial communities in each habitat (Knights et al., 2011; Xiong et al., 2021b). To further support the microbial source analysis, nestedness analysis was performed. The temperature statistics (*T*, smaller the *T* value, perfect the nestedness), based on pairwise compositional difference, and the nestedness metric, based on overlap and decreasing fill, were calculated using the R packages vegan and bipartite (Bernard et al., 2021).

Null model and βNTI (β-nearest taxon index metrics) analyses were calculated using the picante R package for distinguishing different community ecological processes, including deterministic (|βNTI| > 2) and stochastic process (|βNTI| > 2) (Kembel et al., 2010). Specifically, based on the βNTI and the Bray-Curtis-based Raup-Crick (RC_bray_), the two ecological processes were divided into five processes: heterogeneous selection (βNTI < − 2), homogeneous selection (βNTI > +2), dispersal limitation (|βNTI|< 2 and RCBray > 0.95), homogenizing dispersal (|βNTI|< 2 and RCBray < –0.95), and undominated (|βNTI|< 2 and |RCBray|< 0.95) (Tripathi et al., 2018).

## 3. Results

### 3.1. Effects of revegetated shrubs on microbial communities

A total of 48 bacterial phyla were observed in both shrub-associated and bulk soil samples. The bacterial communities were dominated by Proteobacteria, Actinobacteria, Bacteroidetes, Acidobacteria, Chloroflexi, and Planctomycetes (Figure. 1a). Notably, Cyanobacteria and Tenericutes were highly abundant in the *S. psammophila* samples. In the bare sandy land plot, Gemmatimonadetes and Planctomycetes also were the dominant bacterial phyla. Specifically, the shrub microhabitats recruited Fusobacteria, which were not detected in the bulk soil samples. At the family level, *Burkholderiaceae, Chitinophagaceae*, and *Sphingomonadaceae* were the top three families in the bacterial assemblage in shrub samples, while the following taxonomic ranks were dramatically different. In the fungal family, the taxonomic ranks varied markedly across shrub and bare sandy land samples. Moreover, nine fungal phyla were found in all the samples (Figure. 1b). Ascomycota was the most abundant phylum in all the samples, whereas Basidiomycota, Chytridiomycota and unclassified taxa were highly abundant in bulk soil samples. Additionally, Blastocladiomycota only was found in *C. korshinskii* and *S. psammophila* samples.

**Figure 1.**
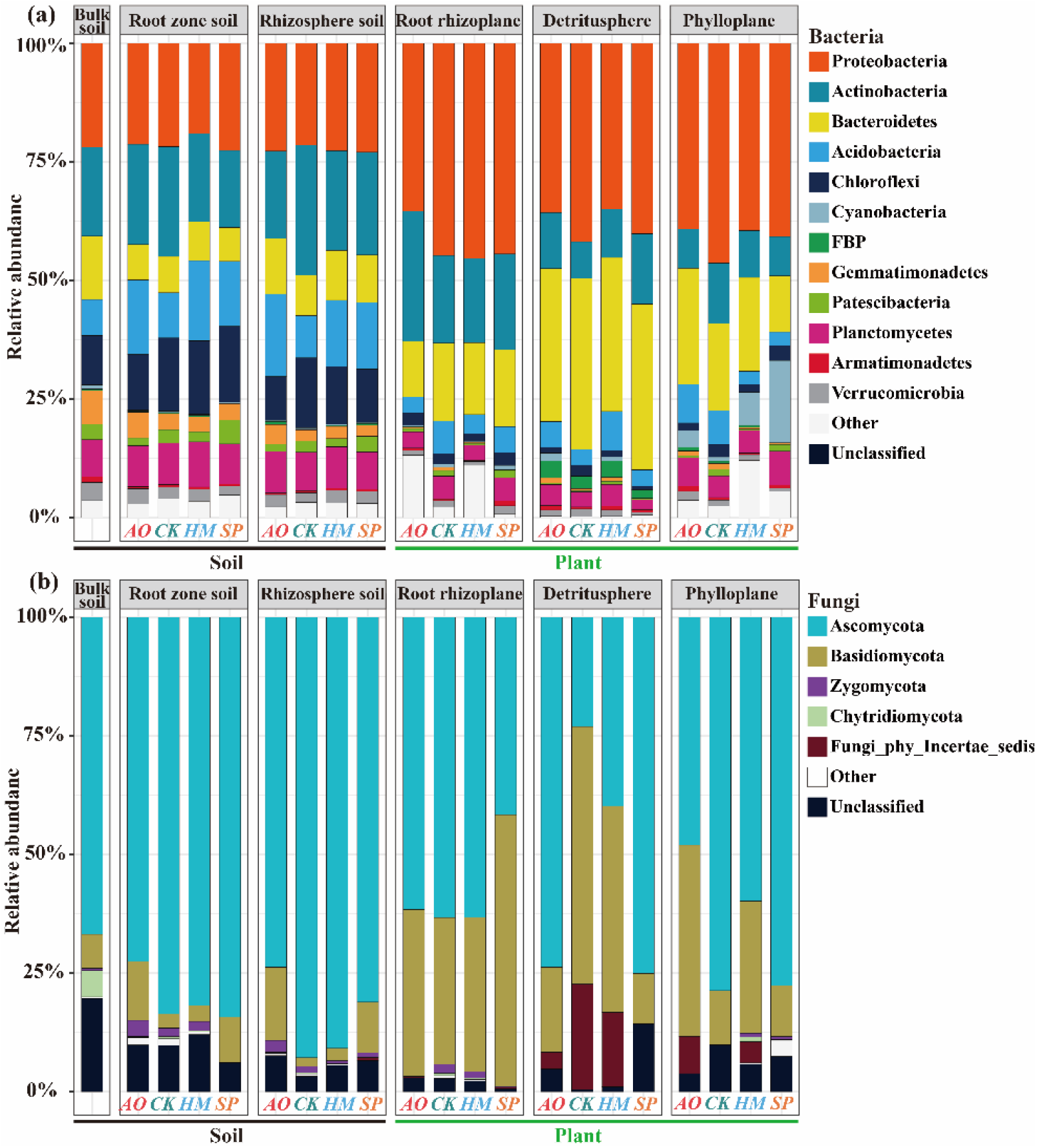
Relative sequence abundance of bacterial (a) and fungal (b) phyla associated with six microhabitats (bulk soil, root zone soil, rhizosphere soil, root rhizoplane, detritusphere, and phylloplane) across four shrublands (*A. ordosica, C. korshinskii, H. mongolicum*, and *S. psammophila*; n = 12) and bare sandy land (n = 12). Operational taxonomic unit with relative abundance < 0.1% were discarded.

The α-diversity of bacteria and fungi (Observed species, Chao1 index, Shannon diversity index, and Goods coverage) were not significantly different across shrub plantations (*p* > 0.05; Figure. S1). However, bacterial Goods coverage in *A. ordosica* samples was lower than that in *S. psammophila* (*p* < 0.01; Figure. S1a). Beta-diversity based on average Bray-Curtis distances was markedly different across four shrub plantations and shrub species explained far greater variation in fungal community composition (Adonis: degree of freedom (d.f.) = 3; coefficient of determination (*R*^2^) = 0.087; *p* < 0.001) than in bacterial community composition (Adonis: d.f. = 3; *R*^2^ = 0.033; *p* < 0.001; Figures. 2a and b; Table 1).

**Table 1.**
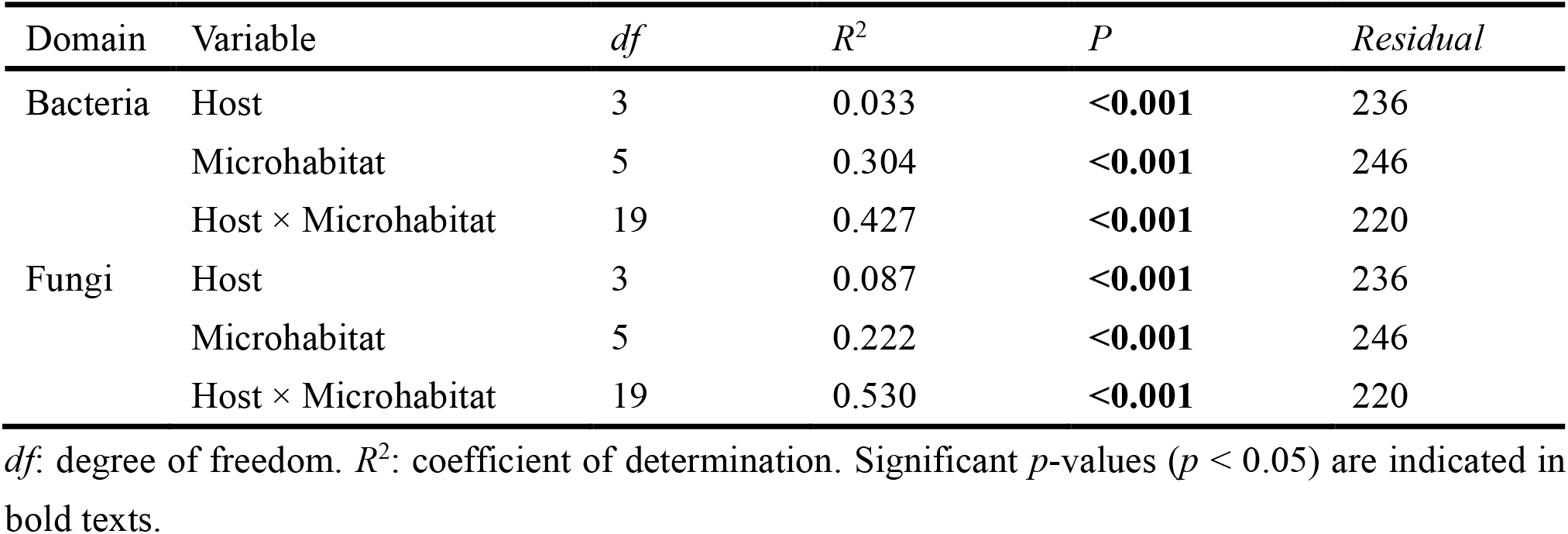
PERMANOVA (Bray-Curtis distance) analysis showing the ability of variables to explain compositional variance.

| Domain | Variable | $df$ | $R^2$ | $P$ | Residual |
| --- | --- | --- | --- | --- | --- |
| Bacteria | Host | 3 | 0.033 | **<0.001** | 236 |
|  | Microhabitat | 5 | 0.304 | **<0.001** | 246 |
| | Host × Microhabitat | 19 | 0.427 | **<0.001** | 220 |
| Fungi | Host | 3 | 0.087 | **<0.001** | 236 |
|  | Microhabitat | 5 | 0.222 | **<0.001** | 246 |
| | Host × Microhabitat | 19 | 0.530 | **<0.001** | 220 |

$df$ : degree of freedom. $R^2$ : coefficient of determination. Significant $p$ -values ( $p < 0.05$ ) are indicated in bold texts.

**Figure 2.**
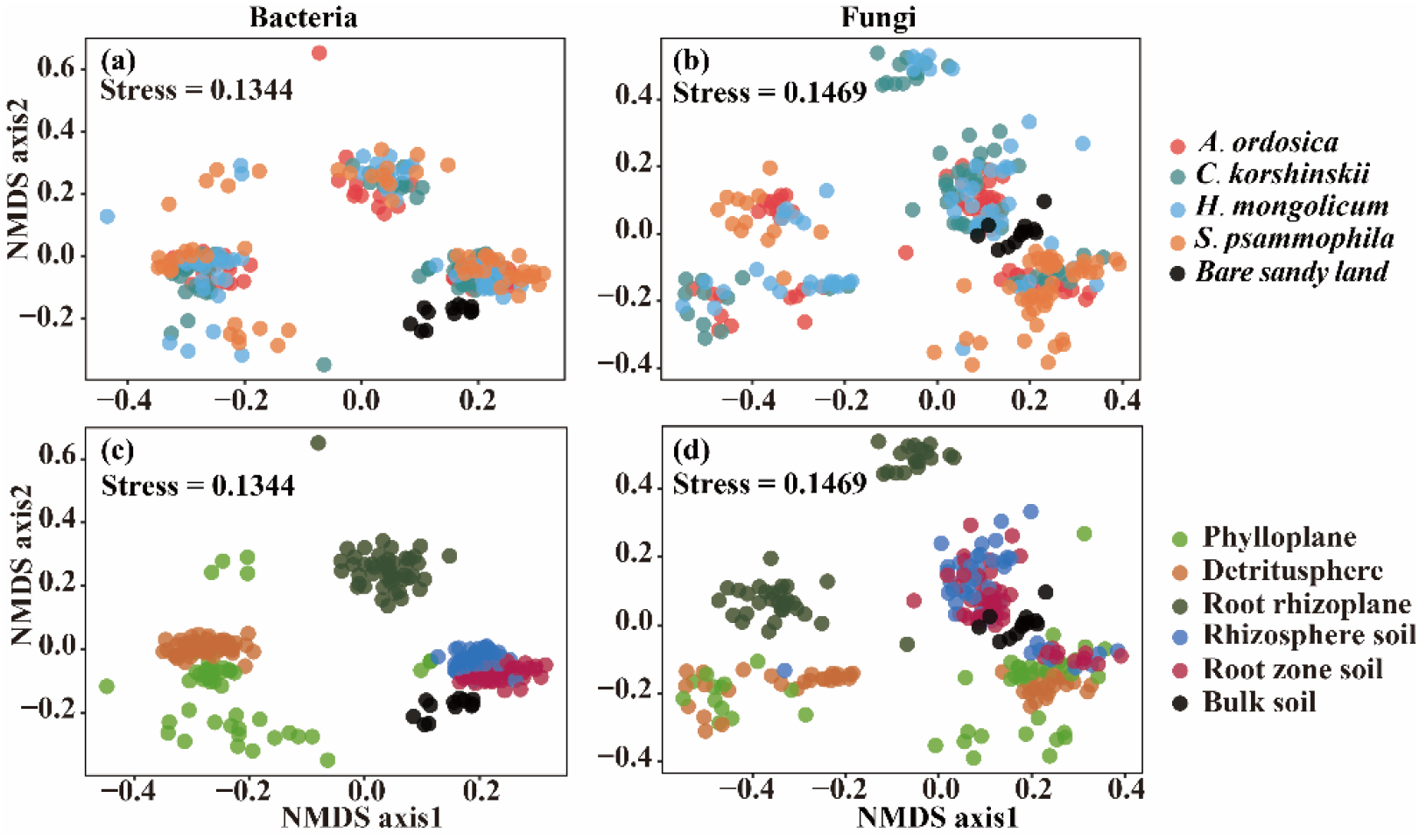
Factors shaping the composition of microbial community of xeric shrubs. (a) and (b) Non-metric multidimensional scaling plots of bacterial and fungal community dissimilarity with shapes by four shrub species (*A. ordosica, C. korshinskii, H. mongolicum*, and *S. psammophila*) and bare sandy land. (c) and (d) Non-metric multidimensional scaling plots of bacterial and fungal community dissimilarity with shapes by six microhabitats (bulk soil, root zone soil, rhizosphere soil, root rhizoplane, detritusphere, and phylloplane).

The DESeq2 differential abundance analysis showed that approximate 5.1% of bacterial OTUs and 4.5% of fungal OTUs, mainly belonged to the bacterial families *Sphingomonadaceae* and *Chitinophagaceae*, and the fungal families *Pezizomycotina_fam_Incertae_sedis* and *Trichocomaceae*, respectively, were significantly enriched in four revegetated shrublands (Figure. 3; Table S2). We also found that approximate 8.2% of bacterial OTUs and 4.5% of fungal OTUs significantly depleted in four revegetated shrublands, these OTUs were mainly from the bacterial families *Gemmataceae* and *Gemmatimonadaceae*, and the fungal family *Pezizomycotina_fam_Incertae_sedis* (Figure. 3; Table S2). Specially, *S. psammophila* possessed the lowest numbers of enriched OTUs for bacteria, while *A. ordosica* possessed the greatest numbers of enriched OTUs for fungi.

**Figure 3.**
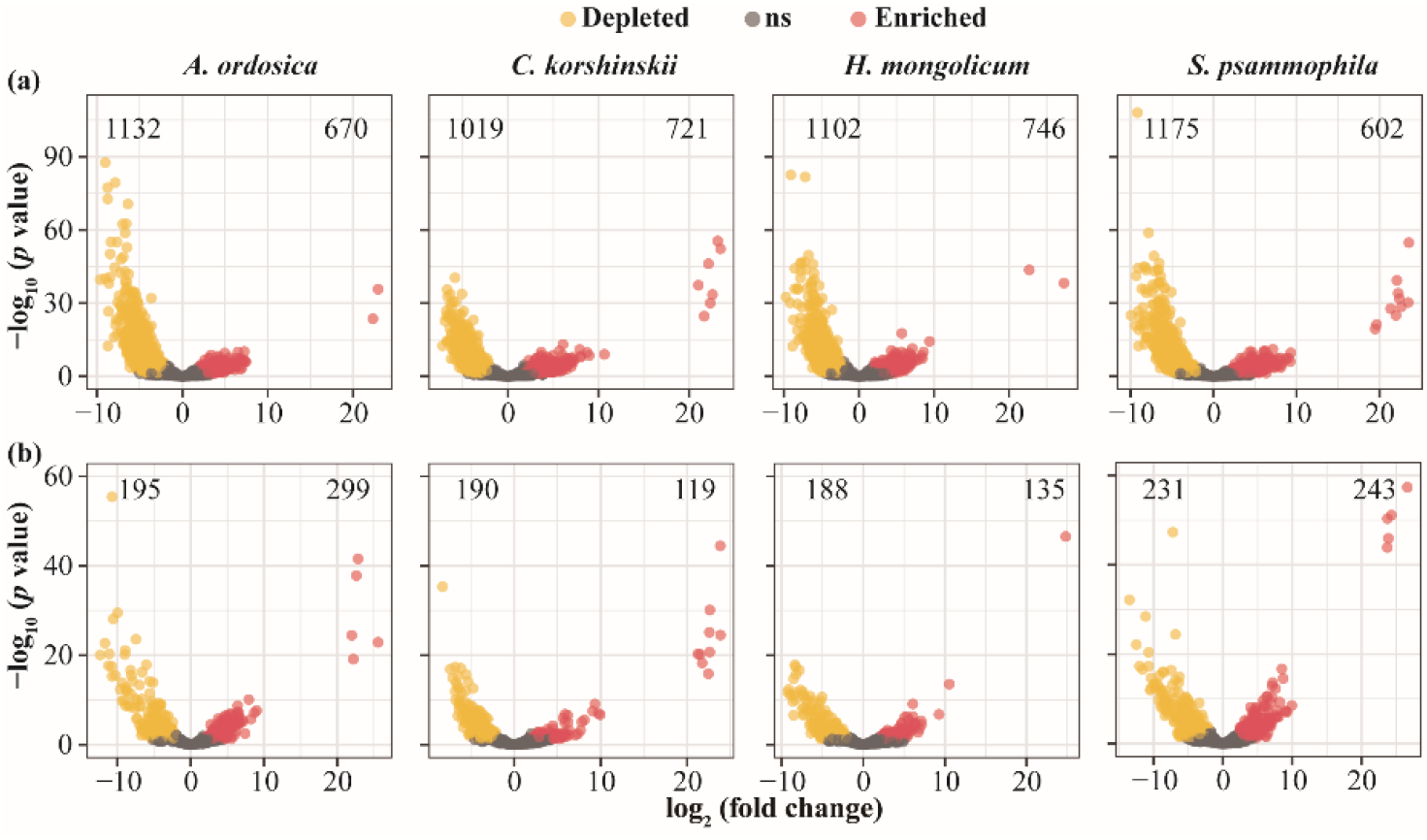
The volcano plot illustrating the enrichment and depletion patterns of the shrub-associated bacterial (a) and fungal (b) communities in different revegetated shrublands (*A. ordosica, C. korshinskii, H. mongolicum*, and *S. psammophila*) compared with bare sandy land.

### 3.2. Niche differentiation of shrub-associated microbiota across different microhabitats

The dominant bacterial and fungal phyla dramatically differed across the microhabitats (phylloplane, detritusphere, root rhizoplane, rhizosphere soil, root zone soil, and bulk soil; *p* < 0.01; Figure. 1). Proteobacteria and Bacteroidetes had higher abundance in plant tissue samples than soil samples, but the abundance of Acidobacteria and Planctomycetes were lower in plant tissue samples (*p* < 0.001). Specifically, Proteobacteria, Bacteroidetes, Actinobacteria, Acidobacteria, and Planctomycetes (average across shrub species; n = 12) were abundant in all microhabitats, whereas Cyanobacteria, FBP, Tenericutes, and Chloroflexi were more abundant in leaves, detritus, roots, and soils, respectively, compared to other microhabitats (Tukey’s honestly significant difference test: *p* < 0.01). Ascomycota and Basidiomycota were the most dominant, accounting for approximately 85% of the sequences. Interestingly, more fungal phyla were detected in the leaf samples than in the other samples. Blastocladiomycota was only detected in leaf samples of *C. korshinskii* and *S. psammophila*.

The Observed species, Chao1 index, Shannon index, and Goods coverage values showed a similar trend for bacterial α-diversity among microhabitats, with a significant difference between the soil microhabitats (rhizosphere, root zone, and bulk soil) and the plant tissue microhabitats (root rhizoplane, detritusphere, and phylloplane) (Kruskal–Wallis test and Dunn’s post-hoc test, *p* < 0.001; Figure. S2). Specially, in the plant tissue microhabitats, the Observed species and Shannon index of the detritusphere were significantly higher than other microhabitats (*p* < 0.05). In the soil microhabitats, the root-associated soil had greater Chao1 index than bulk soil did (*p* < 0.01). For fungi, the α-diversity index had a dramatical difference between detritusphere and other microhabitats, with a slight difference in Observed species and Shannon index (Figure. S2b).

The differential abundance analysis demonstrated that root rhizoplane possessed the greatest numbers of enriched OTUs, mainly from the bacterial family *Gemmataceae* and the fungal family *Pezizomycotina_fam_Incertae_sedis* (Figure. 4; Table S3). Meanwhile, the lowest numbers of depleted OTUs, mainly belonged to the bacterial family *Chitinophagaceae* and the fungal family *Thelephoraceae* was also observed in root rhizoplane (Figure. 4; Table S3). In soil microhabitats (rhizosphere and root zone soil), the enriched OUTs for bacteria and fungi were mainly assigned to the families *Gemmatimonadaceae* and *Gemmataceae*, and *Spizellomycetaceae* and *Lasiosphaeriaceae*, respectively. Compared to soil microhabitats, the aboveground microhabitats (detritusphere and phylloplane) had the greater numbers of the enriched OTUs, mainly from the bacterial families *Gemmatimonadaceae* and *Pirellulaceae*, and the fungal families *Pezizomycotina_fam_Incertae_sedis* and *Trichocomaceae* (Table S3).

**Figure 4.**
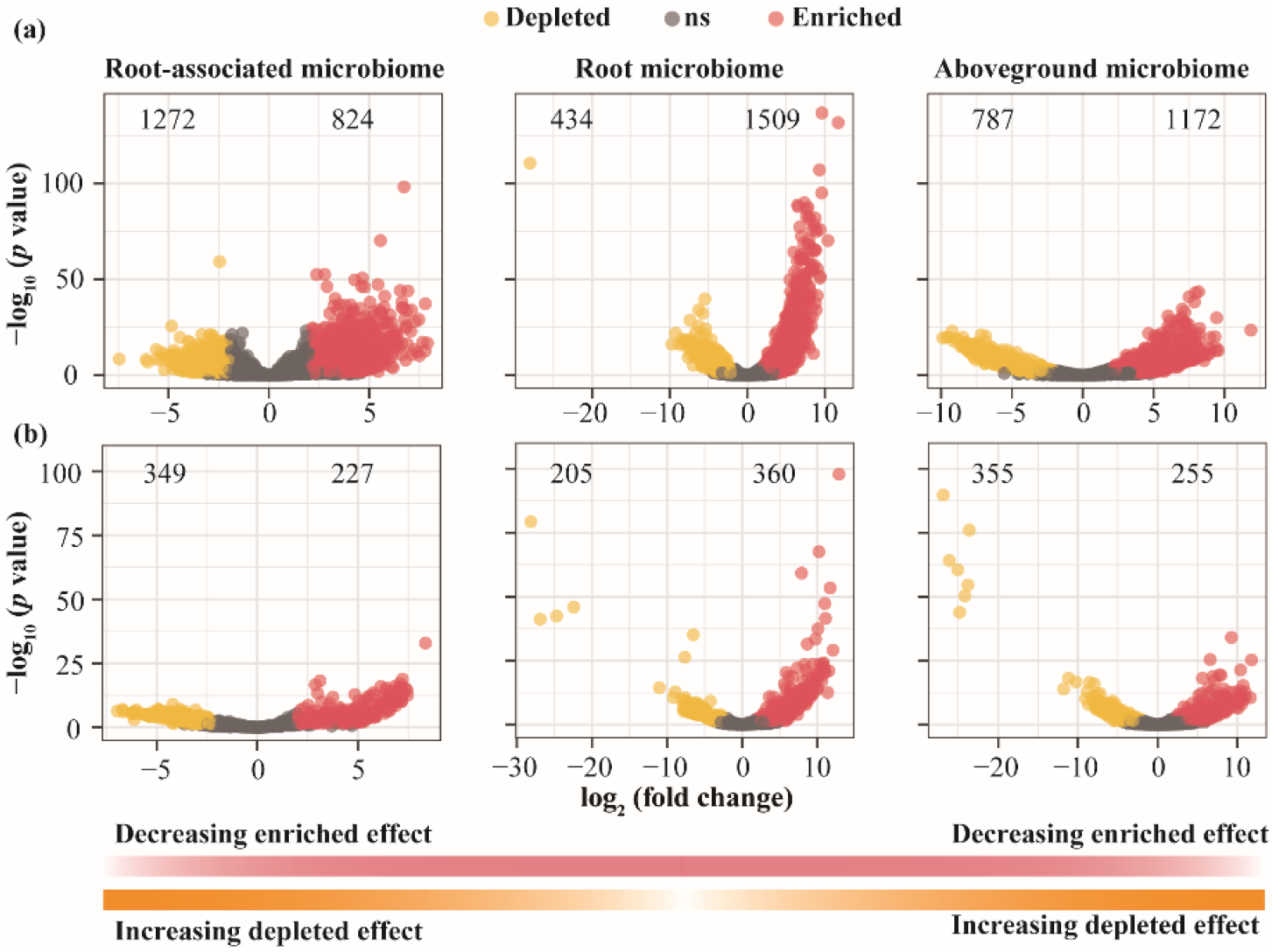
The volcano plot illustrating the enrichment and depletion patterns of the shrub-associated bacterial (**a**) and fungal (**b**) communities in six microhabitats (phylloplane, detritusphere, root rhizoplane, rhizosphere soil, and root zone soil) compared with bulk soil.

Microhabitat was the primary factor explaining the variation in shrub-associated microbial community composition (Adonis: d.f. = 5; bacteria: *R*^2^ = 0.30; *P* < 0.001; fungi: *R*^2^ = 0.22, *p* < 0.001; Table 1). PERMANOVA, conducted separately for shrubs, indicated that the microbial community composition significantly varied among different microhabitats (Table S4; *p* < 0.01). Community similarity analysis showed that rhizosphere, root zone, and bulk soil samples were closely related to each other and detritus samples were the most similar to leaf samples. In addition, root rhizoplane samples were dissimilar to the other samples.

The network analysis showed that microbial co-occurrence patterns differed distinctly across six microhabitats, particularly for the bulk soil and root rhizoplane (Figures. 5a and b, Tables S5 and S6). Bacterial network in bulk soils was the most complex, followed by phylloplane and root-associated microhabitats (root zone soils, root rhizoplane, and rhizosphere soils), with the lowest bacterial network complexity in the detritusphere. For fungi, the highest and lowest network complexity was found in the root rhizoplane and rhizosphere soils, respectively. The network complexity in bulk soils was greater than that in other microhabitats (phylloplane > detritusphere > root zone soils), and the highest modularity and the lowest average path distance were observed in the bulk soil. We defined the “network hubs” (degree > 50; closeness centrality > 0.3) in the network, and found 1038 network hubs (bacteria: 697, fungi: 341) at bulk soil, and 157 network hubs (bacteria: 0, fungi: 157) at root rhizoplane, and 51 network hubs (bacteria: 0, fungi: 51) at detritusphere (Figure. 5c, Table S5). In the bacterial network, a half of nodes were assigned to the top 3 phyla (Proteobacteria, Planctomycetes, Actinobacteria) in soil microhabitats (bulk soil, root zone soil, rhizosphere soil), whereas in plant microhabitats (root rhizoplane, detritusphere, phylloplane) the top-three phyla were Proteobacteria, Bacteroidetes, and Actinobacteria (Figure. 5b, Table S6). For fungi, two phyla (Ascomycota and Basidiomycota) were identified in nodes, accounting for approximate 80% of all nodes. Remarkably, Zygomycota and Glomeromycota were not detected in network nodes of detritusphere (Figure. 5b, Table S6).

**Figure 5.**
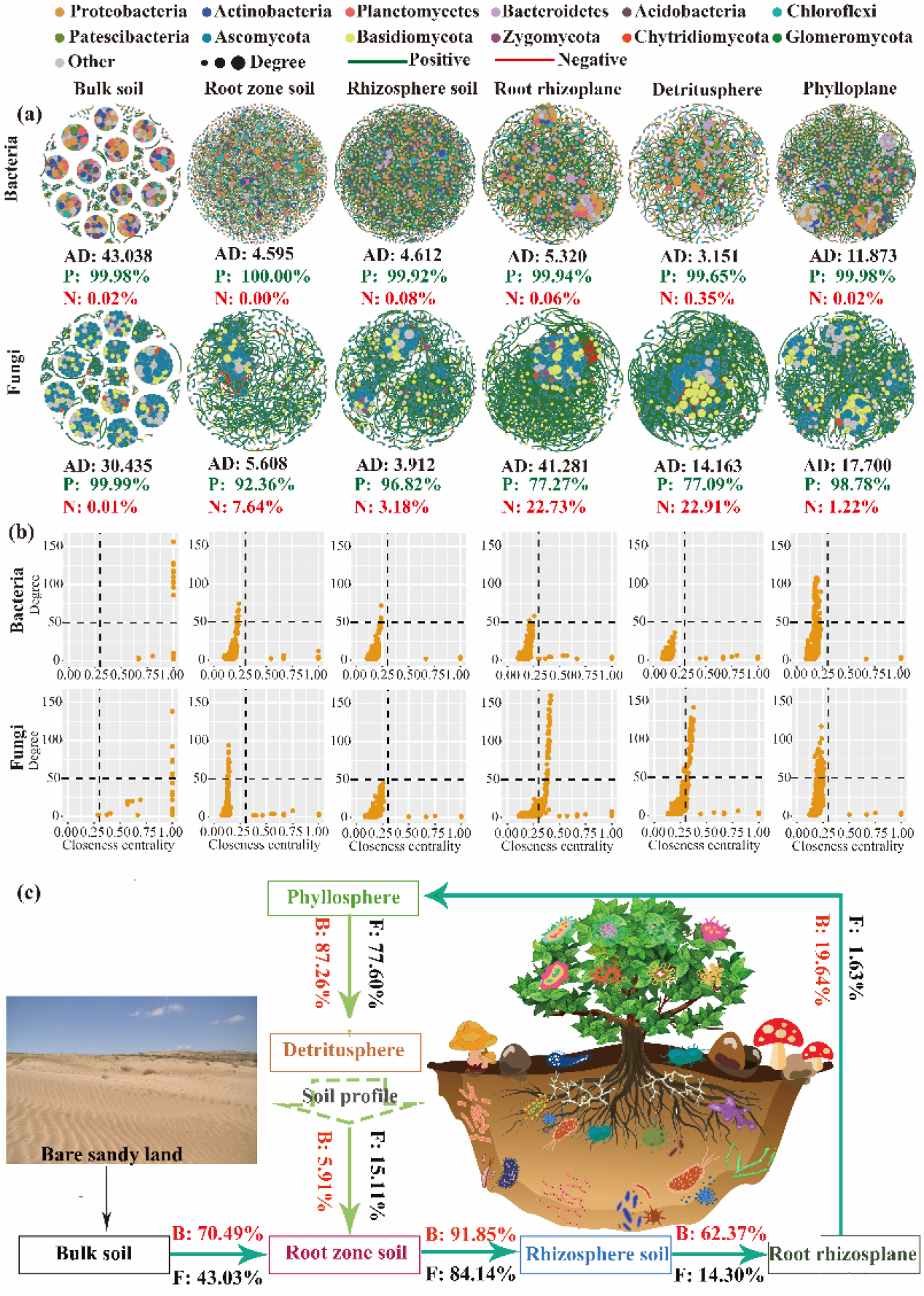
Spatial dynamics of xeric shrub-associated microbiomes. (a) Bacterial and fungal co-occurrence networks along the soil-plant continuum (n = 48). AD: average degree. Different colours represent microbial phyla. (b) Distribution patterns of the hub nodes (degree > 50; closeness centrality > 0.3) of bacterial and fungal network in different microhabitats. (c) Source model analysis based on the SourceTracker showing the potential sources of xeric shrub-associated microbiota (n = 48).

### 3.3. Potential sources of shrub-associated microbiota

The SourceTracker analyses suggested that root-associated bacterial communities were mainly derived from bulk soils and gradually transmitted to different belowground microhabitats (Figure. 5c). Nevertheless, similar source patterns were not observed in root-associated fungal communities. Plant tissue niches accounted for a smaller proportion of derivation of fungal communities than of bacterial communities. Phylloplane, the main potential source of detritusphere, acquired a minority of taxa from the belowground species pool (bacteria: 19.64%, fungi: 1.63%). Conversely, the aboveground species pool was also a potential source of soil microhabitats, specifically, the detritusphere contributed 5−15% of sources to the subterranean microbiotas (Figure. S5). Nestedness analysis also showed that bacterial communities were more perfectly nested by habitats than fungal communities (bacteria: *T* = 9.74°, *P* = 0.01, NODF = 19.28%; fungi: *T* = 31.38°, *P* = 0.01, NODF = 57.61%), with rhizosphere soils having the highest microbial diversity. Specifically, fungal communities in the phylloplane were an important species subset.

### 3.4. Assembly processes of shrub-associated microbiomes

Null model analyses revealed that the relative importance of determinism vs. stochasticity in shrub-associated bacterial and fungal communities varied across six microhabitats (Figure. 6). After quantifying the deviation in βNTI values, we observed that deterministic assembly processes, especially homogeneous selection, represented a predominantly higher percentage than stochastic assembly processes in bacterial communities, while the stochastic assembly processes (dispersal limitation and undominated processes) were dominant in fungal communities (Figure. 6). Notably, for belowground habitats, the relative contribution of determinism and stochasticity showed a slightly increasing tendency from bulk soil to root rhizoplane in bacterial and fungal communities, respectively (Figure. 6a).

**Figure 6.**
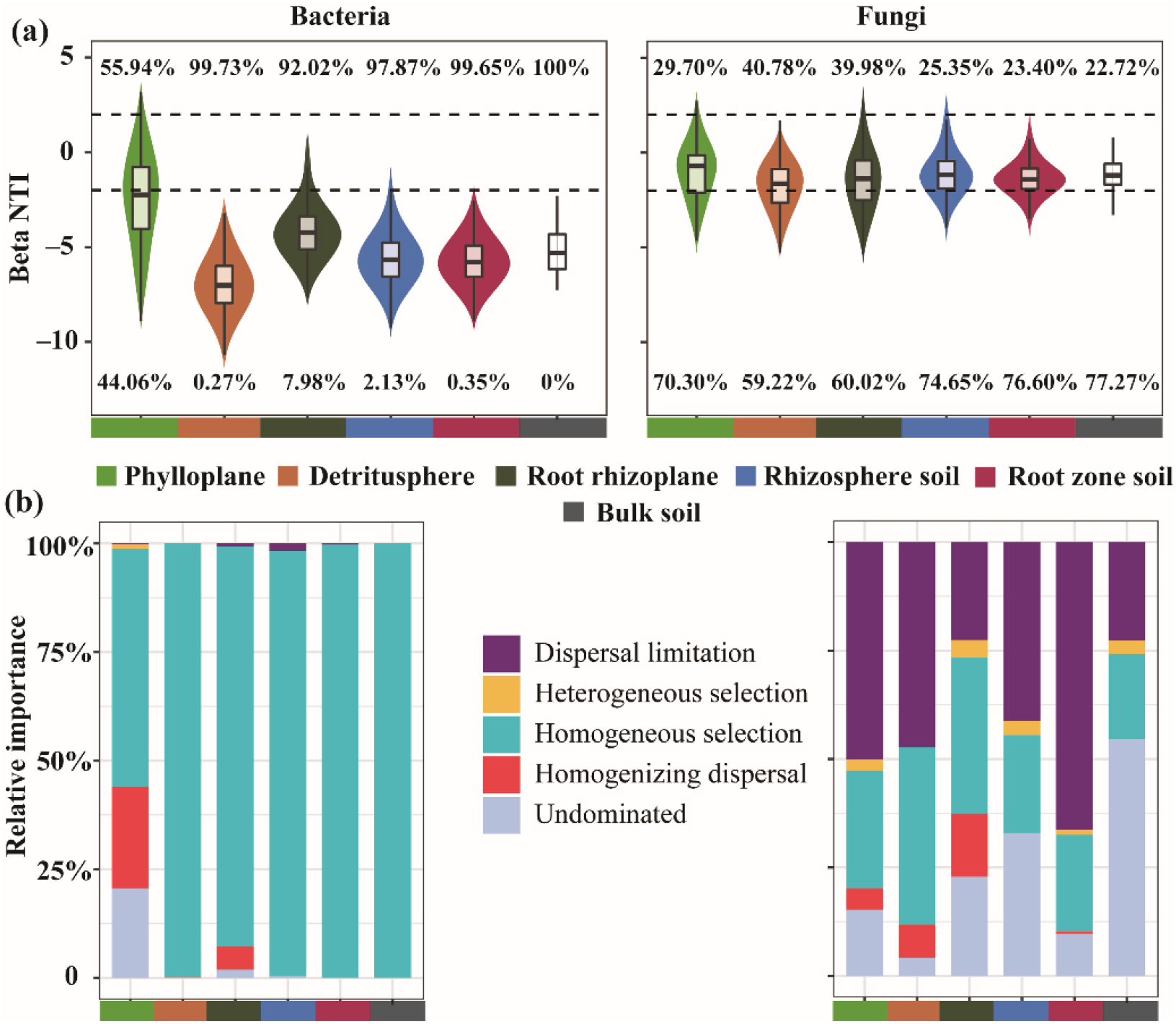
Bacterial and fungal community assembly processes in six microhabitats. (a) The values of the β-Nearest Taxon Index value (βNTI) for shrub-associated microbial communities. Dashed lines represent upper and lower significant thresholds at βNTI = −2 and +2, respectively. (b) The fraction of shrub-associated microbial community assembly governed principally by deterministic processes (heterogeneous selection and homogeneous selection) or stochastic processes (dispersal limitation, homogenizing dispersal, and undominated processes).

## 4. Discussion

### 4.1. Shrubs harbour different microbial communities across different microhabitats

Although the α-diversity analysis indicated that there were no dramatic differences across four shrublands, results of PERMANOVA and differential abundance analysis supported the first two hypotheses that the four revegetated shrub species recruit markedly distinct bacterial and fungal communities after colonisation on bare sandy land, with a great difference in microbial compositions across microhabitats (phylloplane, detritusphere, root rhizoplane, rhizosphere soil, and root zone soil; Figures. 1-4, S1-S4). In addition, niche differentiation, rather than plant species, shaped the diversity and structure of microbiomes (Table 1). Previous studies have shown that the introduction of shrubs on bare sandy land changed the abundance, diversity, and composition of soil bacterial and fungal communities in the degraded dryland ecosystem (Cregger et al., 2018; Sun et al., 2019), whereas our results provide a more precise case assessment of the promotion of biodiversity in revegetated drylands.

In the results, the pattern of niche driving to microbial was supported by some studies on crops (Xiong et al., 2021a, b) and wild plants (Cregger et al., 2018; Zheng and Gong, 2019; Wang et al., 2022) from local to regional areas. However, recent studies have shown that fungal compositional variance is better predicted by sampled sites than by microhabitats in regional areas (Bernard et al., 2021; Thiergart et al., 2020). This phenomenon was confirmed in the present study site where the four shrubs encountered similar growth conditions (soil and climate) but the numerous microenvironment properties (e.g., nutrients, temperature, humidity, and plant immunity) varied remarkably across the soils and plant tissues. Moreover, the microbial co-occurrence network analysis indicated that the different topological structures (i.e., network complexity, modularity, clustering coefficient, and hub node) between soils and plant habitats influenced the bacterial communities more than the fungal communities (Figures. 1 and 5; Tables S5 and S6). Another explanation may be that fungi are more easily affected by environmental conditions due to the existence of some fungal communities not directly associated with a plant (Gao et al., 2020). Additionally, in desert soil with low nutrients, plants supply most of the soil organic matter for microbiomes, consequently resulting in a strong relationship between plants and microbes (Hu et al., 2021). All these evidences suggested that, after revegetation in drylands, the recruitment and domestication to soil microbiome could depend on host selection and microhabitat differentiation caused by different plant species.

Notably, revegetated shrubs had significantly different strategies for microbiomes colonisation (Figures. 1−3, S1, and S3), implying that this process majorly influences the soil health and multifunctionality. Plants create several microhabitats (phylloplane, detritusphere, root rhizoplane, rhizosphere) with drastic environmental conditions for colonisation of the surrounding microbiome (Cordovez et al., 2019; Trivedi et al., 2020). In addition, plants also supply different litters and secretions to attract microbiomes and adapt secretion components according to the environmental changes (Zhalnina et al., 2018). For example, in phosphorus-limited soils, legume roots mainly secrete citric acid, fumaric, malonic, succinic, and malic acids, which are favourable for Proteobacteria growth, whereas in phosphorus-rich soils, there are high levels of exudates of citrate, malate, and oxalate, stimulating growth of Acidobacteria (Dai et al., 2020). Volatile oil from *A. ordosica* influences the growth of desert soil microalgae *Palmellococcus miniatus*, thereby, affecting the surrounding microbiomes (Yang et al., 2012). Thus, the role of plant natural accessions (i.e., root exudate, root litter, leaf litter) on the microbiome assembly should be intensively studied.

The co-occurrence network in the plantations had a lower complexity and clustering coefficient and a higher average path distance than those in bare sandy land, indicating a less compact microbial association in the plant compartments than in bare sandy land (Figures. 5a, b, Tables S5 and S6). Plant inputs (litter and exudates) change the cross-feeding relationships of microbes (Malik et al., 2020), which reduces belowground competition for organic matter (Hu et al., 2021). The encroachment of plants results in changes in the microenvironment (i.e., air temperature, soil erosion, and edaphic properties), which indirectly affects the microbial community and their associations (Hu et al., 2021). In contrast, in bare sandy land with limited nutrition, metabolic exchange likely promotes microbial survival and assembly (Leff et al., 2017). These results imply that the microbial compositions of non-plant and plant land are largely shaped by metabolic interactions and resource competition, respectively. Although our study provides some insights into the mechanisms underlying the plant microbiomes, longitudinal experiments should be conducted (Bai et al., 2020).

In this study, the highest network complexities were found in the phylloplane and root rhizoplane, respectively for bacteria and fungi in shrublands (Figures. 5a, b; Table S5). This could be explained by intense competition for nutrition at the interface (roots and leaves) between the plant and the environment (Mommer et al., 2016; Remus-Emsermann and Schlechter, 2018). Broad ecological differences in substrate preferences, growth rates, and stress tolerance lead to distinct trajectories in bacteria and fungi during plant recovery (Sun et al., 2017). For example, fungi are generally considered as the major decomposers of recalcitrant organic matter because of their ability to produce specific enzymes (Bani et al., 2018). Additionally, mycorrhizal fungi can directly clone living plant tissues (Tedersoo et al., 2014). Hyphae filamentous fungi also provide ecological opportunities for bacteria, leading to novel host-symbiont interactions (Emmett et al., 2021; Pawlowska et al., 2018; Yuan et al., 2021). In addition, microbial spores can diffuse via attachment to motile soil bacteria (Muok et al., 2021). In summary, under harsh conditions in drylands, soil microbiomes were extraordinarily sensitive to organic matter input and microenvironment changes via revegetation. However, the response of interspecific and intraspecific interactions between microbiomes (i.e., bacteria and fungi) should have an in-depth exploration.

### 4.2. Source and sink of shrub-associated microbiota

Source-tracking and nestedness analysis showed that a legacy effect of the original land use on shrub-associated microbial assembly and revealed that the legacy effect of bare sandy soil on shrub - associated bacterial communities was stronger than that on shrub-associated fungal communities (Figure. 5c). However, the results partially support the source-sink hypothesis. In concordance with previous studies (Amend et al., 2019; Bernard et al., 2021), rhizospheric soil, not the root zone soil or bulk soil, was identified as the main species reservoir of plant-associated microbial taxa (Figure. S5), further indicating that the rhizosphere act as a hotspot of plant-microbe-soil interactions. However, another study found that plant bacterial communities are gradually filtered and enriched from bulk soils to plant niches (Xiong et al., 2021b). This could be primarily attributed to the harsher soil conditions (low nutrients and soil moisture) in the root zone and bulk soil than in the rhizosphere in the desert ecosystem. In nutrient-poor soils, the rhizosphere, being a hotspot for intense plant-soil-microbe interactions, provides a more pleasant habitat for different microbiomes than that of other soil zones (Mommer et al., 2016), since roots can change the microhabitat environments via dead litter and bioactive exudates (Hu et al., 2018). Thus, our current results, consistent with the previous investigations, show that soil microbial communities under shrub canopies and between shrub canopies have no significant difference (Sun et al., 2019). These results indicate that revegetated plants could prune the original soil microbial communities and modify soil microbiota composition in the whole shrubland. However, future research should focus on the role of shrub root traits (i.e., elongation and turnover) and soil animals (i.e., ants and nematodes) in this process.

Interestingly, in the present study, aboveground plant species pools also contributed to the belowground microbial communities, although microbial diversity of soils was higher than that of plant tissue. In particular, for fungal communities, phylloplane was the second largest species pool (Figure. S5b). These vertically stratified microbiota assembly patterns have also been determined in previous studies in other plant species (Amend et al., 2019). Plants specifically recruit and elaborately prune a small group of beneficial microbes from the soil pool during their lifetime (van der Heijden and Schlaeppi, 2015). A considerable part of plant microbiome diversity, which affects germination and seedling development, may be inherited from the seed (Walsh et al., 2021). Furthermore, experimental evidence indicates that root and phyllosphere microbes are partially inherited via vertical seed transmission (Abdelfattah et al., 2021). In clonal plants, vertical transmission between plant generations occurs in a significant proportion of symbiotic bacteria and fungi (Vannier et al., 2018). Additionally, the air microbiome contributes to phyllosphere microbiota assembly (Archer et al., 2019), which further affects the soil microbiome through rainfall and eluviation. Several fungal spores can disperse onto leaves of neighbouring plants via rain splash, even when wind flow is very low (Mukherjee et al., 2021). An alternative explanation may be that exotic herbivorous insects alter the leaf microbiome through eating leaves and carrying microorganisms, thereby, affecting the soil microbial community via litter input (Humphrey and Whiteman, 2020). Overall, the horizontal and vertical transmission pathways mostly explain the origin and dispersion of microbiomes in plants. However, other mechanisms, such as the effects of leaf-derived microbiomes on the soil microbial community and the contributions of deep soil microbiomes to plant microbiota, warrant further validation.

### 4.3. Ecological assembly processes of shrub-associated microbiota

Disentangling the assembly mechanisms of plant-associated microbiomes is imperative for better understanding the role of plant in generating and maintaining microbial diversity (Trivedi et al., 2020). In this present study, quantitative analysis of assembly processes showed that bacterial and fungal communities across different microhabitats were mainly drove by determinism and stochasticity, respectively (Figure. 6a), partially in contrast to the finding of Cao et al. (2022), who detected that the stochastic processes were dominant in bacterial communities of shrublands in eastern of the Mu Us Desert. This discrepancy can be credited to the difference in precipitations. In drylands, previous studies have proven that precipitation primarily regulated microbial assembly processes, especially bacterial communities (Jiao et al., 2021; Naidoo et al., 2022; Yang et al., 2022), because wetter habitats promote dispersal (Cermeño and Falkowski, 2009). In the current study, bacterial community assembly was dominantly governed by homogeneous selection of deterministic processes (Figure. 6b), indicating that bacteria across six microhabitats had more similar composition (Hanson et al., 2012; Su et al., 2020). Compared to bacterial communities, fungi communities at multiple microhabitats are predominantly govern by dispersal limitation (Bonito et al. 2014; Richter-Heitmann et al., 2020; Xu et al., 2021). Our results supported this view, the proportion of dispersal limitation in fungi at all microhabitats was higher than that in bacteria (Figure. 6b), suggesting that fungi are more limited by resource availability and are more sensitive to environmental changes than bacteria do.

## 5. Conclusions

Our results demonstrate that plant introduction has a much stronger influence on microbial α-diversity and networks in soil microhabitats than plant microhabitats, but the effect on microbial community structure was stronger in plant tissue microhabitats than in soil microhabitats. The changes due to host effect on shrub-associated microbiome composition was stronger at the niche differentiation level rather than at the plant species level in revegetated desert shrubland. We further found that plant microbiome assembly was mainly influenced by plant select and niche filter, meanwhile, revegetated plant via microenvironment changes and microbial seedbank from parent affect soil microbial composition. Furthermore, the surrounding zone of roots is a hotpot for microbial recruitments of revegetated shrubs. Determinism played a dramatically greater role in bacterial communities than fungal communities. Of the four shrubs, *A. ordosica* exhibited the highest performance in plasticity or responsiveness of microbial communities after revegetation; thus, this shrub species is the optimal choice for increasing ecosystem biodiversity in future dryland restoration. Together these results suggest that the host selection (plant niches and host genetics) and soil domestication (organic matter and microbial species seed input) drive microbial community composition and functions in revegetated ecosystems. Collectively, these findings significantly promote our fundamental understanding of the interactions between revegetated plants and microbiomes in drylands during plant introduction. In future studies, the role of plant-associated microbiomes in improving soil nutrient cycle and soil-forming processes of restoring ecosystems should be investigated in depth.

## Acknowledgments

We would like to thank Dr. Chun Miao, Dr. Liang Liu, and Mr. Shijun Liu for their cooperation and assistance in experimental design and field sampling, and the staff of Guangdong Magigene Biotechnology Co., Ltd., Guangzhou, China for their generous support in laboratory work. We also thank Prof. Manuel Delgado-Baquerizo for his constructive and valuable comments and suggestions that helped us improve this article. This work was financially supported by the National Natural Science Foundation of China (Grant No. 31800611 and 31870710) and the National Key Research and Development Plan (No. 2022YFE0104700). Research on soil microbiomes in Yanchi Research Station is supported by the Fundamental Research Funds for the Central Universities (Grant No. PTYX202122 and PTYX202123).

## Author contributions

Zongrui Lai and Yang Yu conceived the research idea. Zongrui Lai, Yanfei Sun, Zhen Liu, Yuxuan Bai, and Shugao Qin sampled in field. Zongrui Lai, Yang Yu, Lin Miao, and Yangui Qiao analysed the data. Zongrui Lai and Yanfei Sun participated in the preparation of this manuscript. Zongrui Lai, Yanfei Sun, and Wei Feng made the illustrations.

## Declaration of Competing Interest

The authors declare that they have no competing financial interests or personal relationships that could have appeared to influence the work reported in this paper.

## Data availability

All RNA-seq data were uploaded to the NCBI Sequence Read Archive (SRA) with the accession number SRP348383.

## Notes

### Competing Interest Statement

The authors have declared no competing interest.

