## Supplementary material for "Plant selection and ecological microhabitat drive domestications of shrub-associated microbiomes in a revegetated shrub ecosystem": Supplementary Material.docx

Address: No. 35 Qinghua Eastroad, School of Soil and Water Conservation, Beijing Forestry University, Haidian District, Beijing 100083, P. R. China

**Supplementary figures**

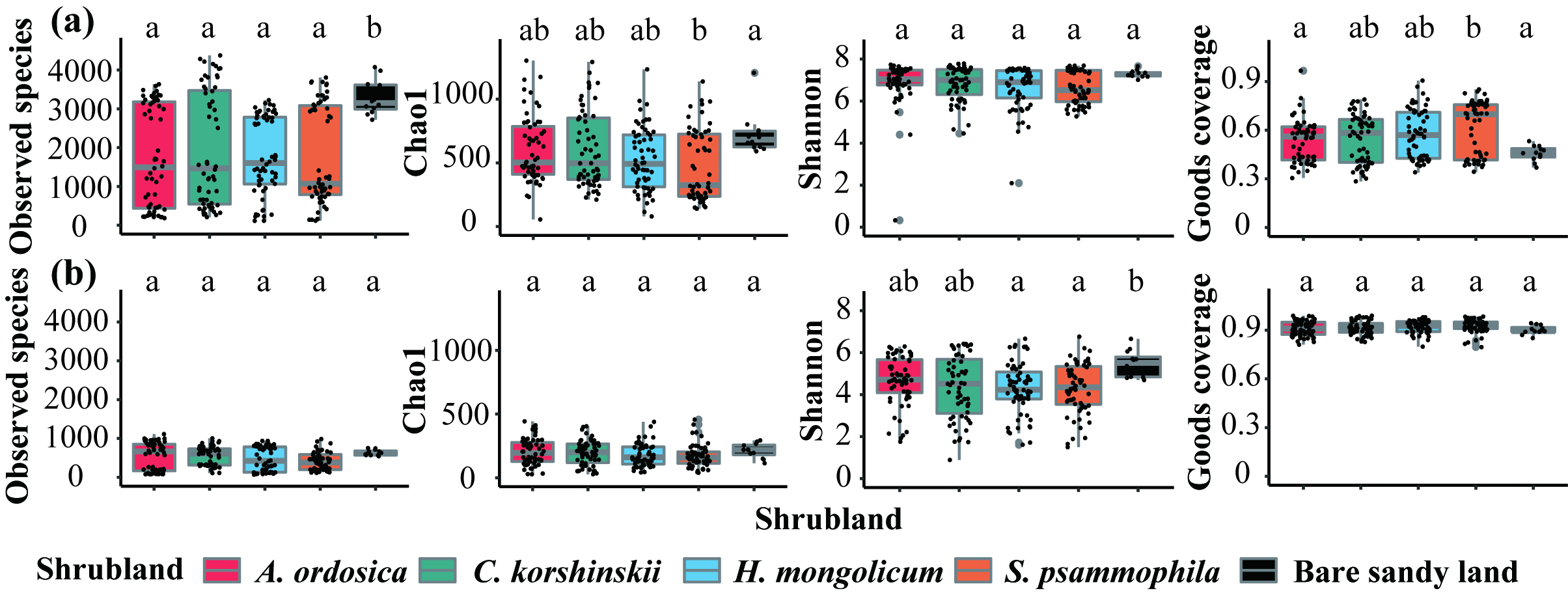

**Figure S1.** The α-diversity, including Observed species, Chao1 index, Shannon index, and Goods coverage values, of bacterial (a) and fungal (b) communities among different revegetated shrublands (A. ordosica, C. korshinskii, H. mongolicum, and S. psammophila, n = 60; bulk soil; n = 12).

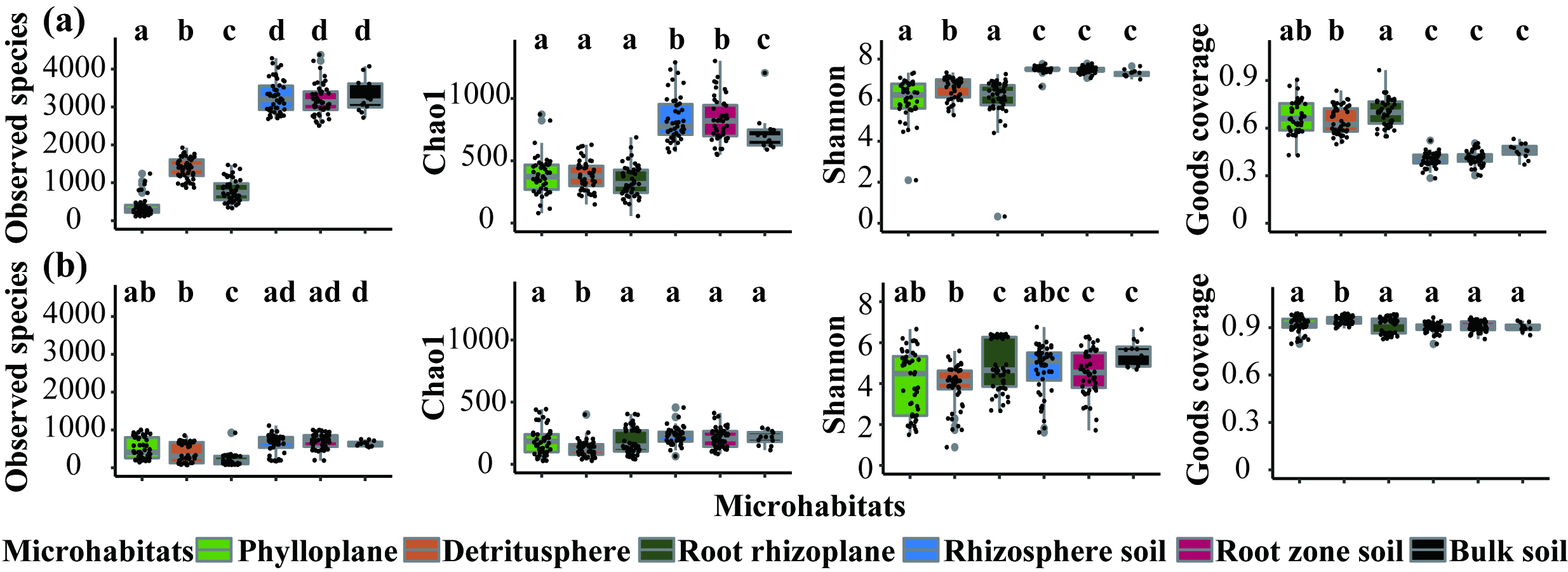

**Figure S2.** The α-diversity, including Observed species, Chao1 index, Shannon index, and Goods coverage values, of bacterial (a) and fungal (b) communities among six microhabitats (n = 48 for phylloplane, detritusphere, root rhizoplane, rhizosphere soil, and root zone soil; n = 12 for bulk soil).

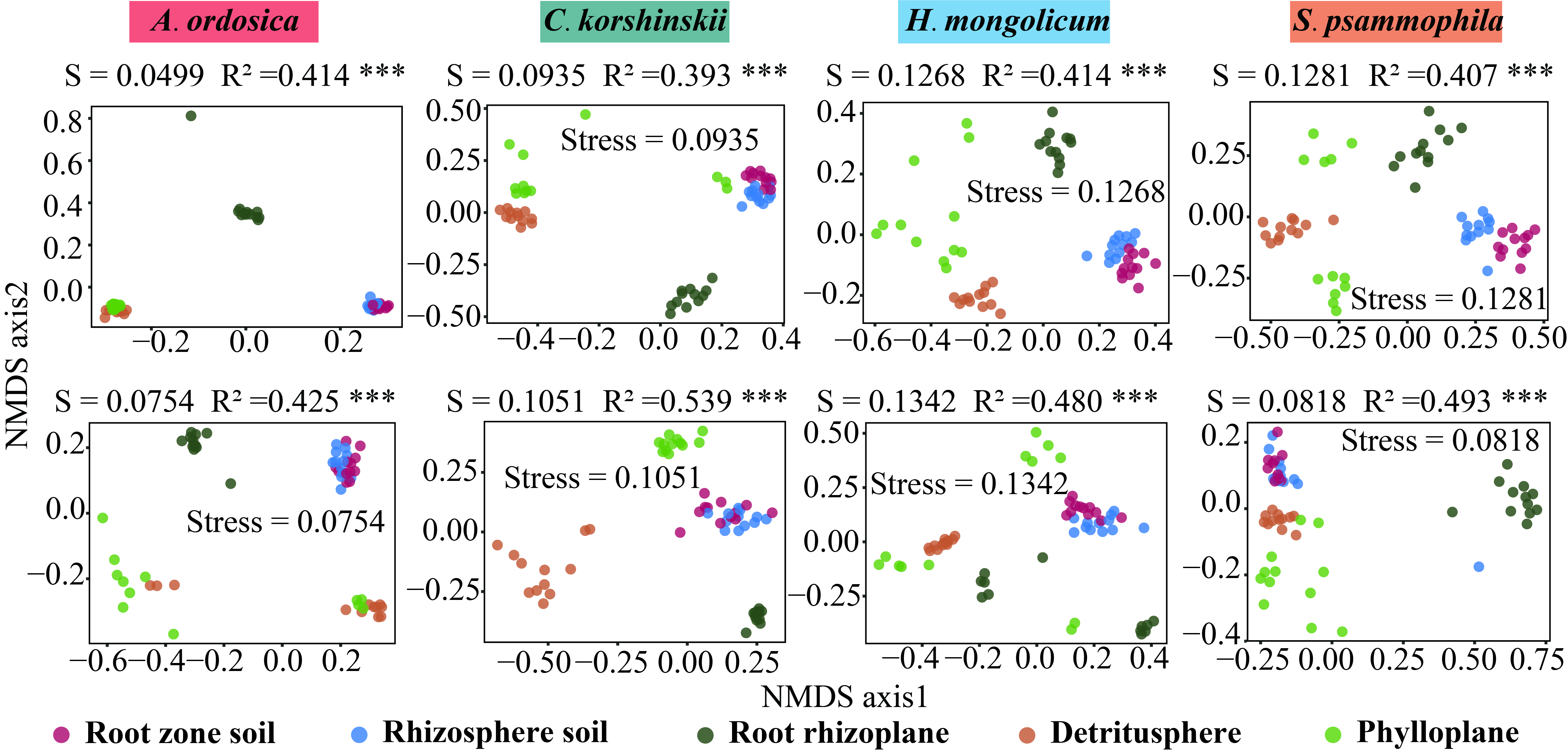

**Figure S3.** NMDS ordination of both bacterial and fungal communities among all the microhabitats (phylloplane, detritusphere, root rhizoplane, rhizosphere soil, and root zone soil) at OUT level for each shrub species (*A. ordosica*, *C. korshinskii*, *H. mongolicum*, and *S. psammophila*)

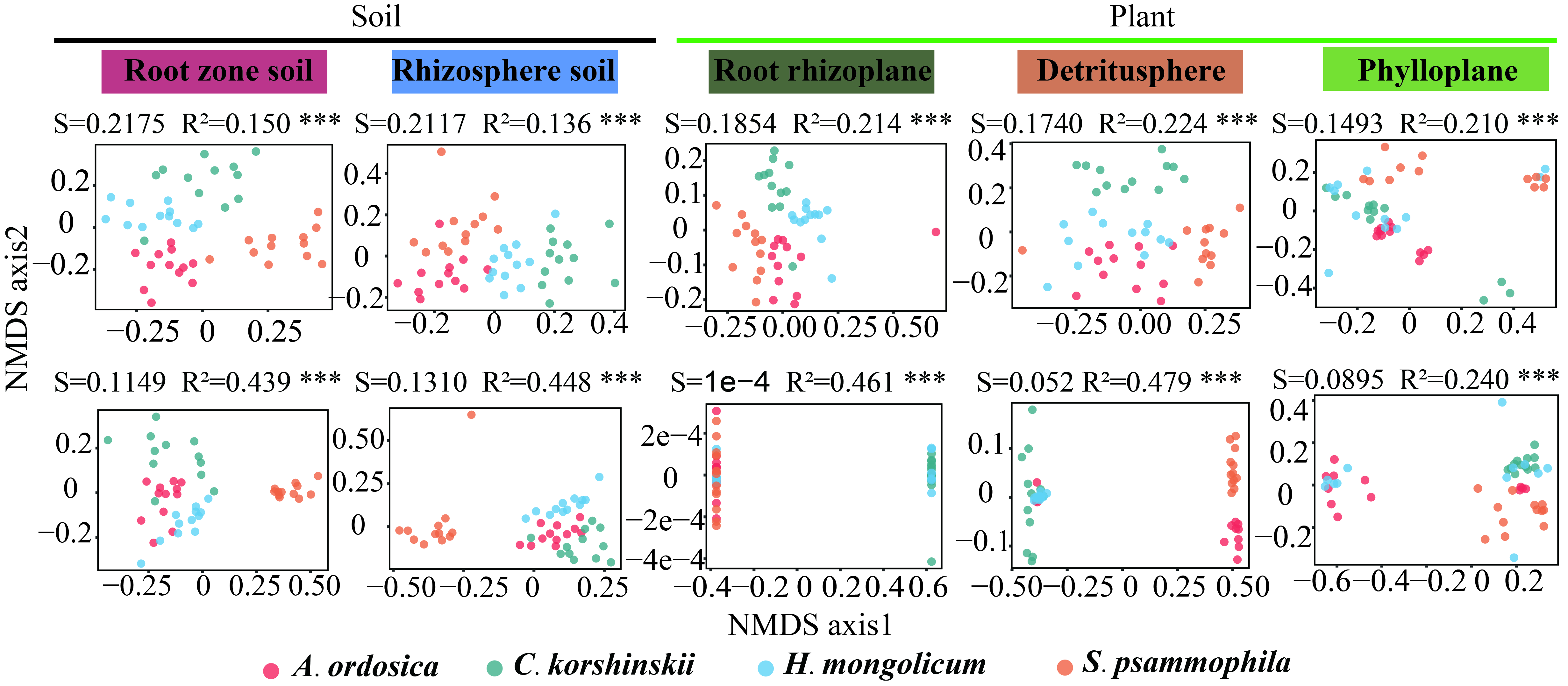

**Figure S4.** NMDS ordination of both bacterial and fungal communities among all the microhabitats (phylloplane, detritusphere, root rhizoplane, rhizosphere soil, and root zone soil) at OUT level for each shrub species (*A. ordosica*, *C. korshinskii*, *H. mongolicum*, and *S. psammophila*)

**Figure S5.** Nestedness plots of overall microbial composition aggregated by microhabitat type. Each vertical line represents an OUT’s presence among microhabitats.

**Supplementary tables**

**Table S1** The soil properties in five study sites.

| Sample plot | SOC  (g·kg^-1^) | TN  (g·kg^-1^) | TP  (g·kg^-1^) | TK  (g·kg^-1^) | NH_4_^+^-N  (mg·kg^-1^) | NO_3_^-^-N  (mg·kg^-1^) | AP  (mg·kg^-1^) | AK  (mg·kg^-1^) | MBC (mg·kg^-1^) | MBN (mg·kg^-1^) | pH | ST  (°C) | SWC  (%) |
| --- | --- | --- | --- | --- | --- | --- | --- | --- | --- | --- | --- | --- | --- |
| *A. ordosica*  *C*. *korshinskii*  *H*. *mongolicum*  *S*. *psammophila*  Bare sandy land | 0.95±0.16  0.60±0.08  0.68±0.11  0.96±0.28  0.67±0.18 | 0.15±0.07  0.20±0.04  0.11±0.02  0.20±0.03  0.12±0.01 | 0.23±0.02  0.20±0.02  0.18±0.01  0.23±0.03  0.19±0.01 | 15.22±0.58  14.39±0.42  14.86±0.79  15.35±0.60  14.70±0.57 | 1.02±0.14  0.91±0.25  0.91±0.15  0.98±0.31  0.30±0.08 | 0.68±0.19  1.02±0.59  1.19±0.51  0.61±0.12  0.38±0.08 | 1.18±0.38  1.63±0.43  1.45±0.25  0.57±0.34  2.13±0.28 | 72.19±36.89  58.40±11.75  82.17±12.55  105.68±33.61  61.60±3.15 | 3.75±2.19  8.82±1.41  5.90±1.91  6.11±2.85  3.14±2.04 | 1.07±0.38  1.28±0.45  0.92±0.41  1.08±0.28  1.29±0.39 | 7.70  7.61  7.66  7.63  7.70 | 25.3  25.1  25.1  24.9  25.7 | 8.5  7.9  8.6  8.9  7.4 |

Soil samples for properties were collected on August 22−23, 2018 (n = 6). The value represents mean ± SD. SOC soil organic carbon, TN total nitrogen, TP total phosphorus, TK total kalium, AP available phosphorus, AK available kalium, MBC microbial biomass carbon, MBN microbial biomass nitrogen, ST sampling soil temperature (light/dark), SWC sampling soil water content.

| Dominant | Shrublands | Depleted | Enriched |
| --- | --- | --- | --- |
| Bacteria  Fungi | *AO*  *CK*  *HM*  *SP*  *AO*  *CM*  *HM*  *SP* | *Gemmataceae* (93)  *Gemmatimonadaceae* (58)  *Pedosphaeraceae* (34)  Unidentified (455)  *Gemmataceae* (70)  *Gemmatimonadaceae* (62)  *Pedosphaeraceae* (32)  Unidentified (412)  *Gemmataceae* (80)  *Gemmatimonadaceae* (63)  *Pedosphaeraceae* (32)  Unidentified (447)  *Gemmataceae* (91)  *Gemmatimonadaceae* (61)  *Pedosphaeraceae* (37)  Unidentified (439)  *Pezizomycotina_fam_Incertae_sedis* (11)  *Glomeraceae* (7)  *Nectriaceae* (4)  Unidentified (88)  *Pezizomycotina_fam_Incertae_sedis* (7)  *Pleosporales_fam_Incertae_sedis* (6)  *Mortierellaceae* (5)  Unidentified (90)  *Pezizomycotina_fam_Incertae_sedis* (10)  *Trichocomaceae* (8)  *Spizellomycetaceae* (4)  Unidentified (81)  *Pezizomycotina_fam_Incertae_sedis* (15)  *Nectriaceae* (8)  *Pleosporales_fam_Incertae_sedis* (7)  Unidentified (99) | *Sphingomonadaceae* (44)  *Chitinophagaceae* (43)  *WD2101_soil_group* (32)  Unidentified (120)  *Chitinophagaceae* (47)  *Sphingomonadaceae* (46)  *Sphingobacteriaceae* (26)  Unidentified (141)  *Chitinophagaceae* (57)  *Sphingomonadaceae* (52)  *WD2101_soil_group* (21)  Unidentified (127)  *Sphingomonadaceae* (39)  *Chitinophagaceae* (30)  *Microscillaceae* (24)  Unidentified (128)  *Pezizomycotina_fam_Incertae_sedis* (17)  *Trichocomaceae* (14)  *Lasiosphaeriaceae* (10)  Unidentified (127)  *Pezizomycotina_fam_Incertae_sedis* (6)  *Trichocomaceae* (5)  *Glomeraceae* (4)  Unidentified (52)  *Chaetomiaceae* (4)  *Inocybaceae* (4)  *Pezizomycotina_fam_Incertae_sedis* (4)  Unidentified (55)  *Pleosporales_fam_Incertae_sedis* (10)  *Tremellales_fam_Incertae_sedis* (6)  *Leptosphaeriaceae* (5)  Unidentified (113) |

**Table S2** Top 3 depleted and enriched families reported by DESeq2 pairwise comparison analysis for bacteria and fungi in each shrubland compared with bare sandy land.

Note: *AO* *A. ordosica*; *CK* *C*. *korshinskii*; *HM* *H*. *mongolicum*; *SP* *S*. *psammophila*.

| Dominant | Microhabitat | Depleted | Enriched |
| --- | --- | --- | --- |
| **Bacteria**  **Fungi** | Soil  Root  Aboveground  Soil  Root  Aboveground | *Gemmataceae* (62)  *Chitinophagaceae* (61)  *WD2101_soil_group* (36)  Unidentified (425)  *Chitinophagaceae* (24)  *Microscillaceae* (17)  *Sphingomonadaceae* (15)  Unidentified (76)  *Sphingomonadaceae* (78)  *Chitinophagaceae* (52)  *Sphingobacteriaceae* (37)  Unidentified (102)  *Pezizomycotina_fam_Incertae_sedis* (15)  *Glomeraceae* (11)  *Agaricaceae* (9)  Unidentified (167)  *Thelephoraceae* (5)  *Mortierellaceae* (4)  *Pyronemataceae* (4)  Unidentified (113)  *Pleosporales_fam_Incertae_sedis* (13)  *Thelephoraceae* (9)  *Orbiliaceae* (8)  Unidentified (167) | *Gemmatimonadaceae* (47)  *Gemmataceae* (46)  *Pedosphaeraceae* (29)  *Unidentified* (305)  *Gemmataceae* (102)  *Gemmatimonadaceae* (69)  *Pirellulaceae* (51)  *Unidentified* (583)  *Gemmataceae* (76)  *Gemmatimonadaceae* (60)  *Pirellulaceae* (41)  Unidentified (102)  *Spizellomycetaceae* (8)  *Lasiosphaeriaceae* (6)  *Pleosporaceae* (4)  Unidentified (147)  *Pezizomycotina_fam_Incertae_sedis* (13)  *Nectriaceae* (8)  *Trichocomaceae* (8)  Unidentified (223)  *Pezizomycotina_fam_Incertae_sedis* (10)  *Trichocomaceae* (9)  *Nectriaceae* (7)  Unidentified (164) |

**Table S3** Top 3 depleted and enriched families reported by DESeq2 pairwise comparison analysis for bacteria and fungi along soil-plant continuum compared with bulk soil.

Note: “Soil” includes rhizosphere soil and root zone soil; “Root” indicates root rhizoplane; “Aboveground” includes detritusphere and phylloplane.

**Table S4** PERMANOVA (Bray-Curtis distance) and pairwise comparisons of microbial community differences among all the microhabitats (phylloplane, detritusphere, root rhizoplane, rhizosphere soil, and root zone soil) at two phylogeny levels.

|  | | Phylogenetic level  pairwise.adonis output | | Bacteria | | | | | | | | | |  | | Fungi | | | | | | | | | |
| --- | --- | --- | --- | --- | --- | --- | --- | --- | --- | --- | --- | --- | --- | --- | --- | --- | --- | --- | --- | --- | --- | --- | --- | --- | --- |
|  |  |  |  | OTU level | | | |  | | Family level | | | |  | | OTU level | | | |  | | Family level | | | |
|  |  | |  | | R^2^ | | *p* | |  | | R^2^ | | *p* | |  | | R^2^ | | *p* | |  | | R^2^ | | *p* |
| *AO* | | Phylloplane vs. Detritusphere | | 0.229 | | *** | | | | 0.438 | | *** | | | | 0.118 | | 0.023* | | | | 0.120 | | 0.042* | |
|  | | Phylloplane vs. Root rhizoplane | | 0.325 | | *** | | | | 0.495 | | *** | | | | 0.328 | | *** | | | | 0.304 | | *** | |
|  | | Phylloplane vs. Rhizosphere soil | | 0.375 | | *** | | | | 0.780 | | *** | | | | 0.273 | | *** | | | | 0.329 | | *** | |
|  | | Phylloplane vs. Root zone soil | | 0.386 | | *** | | | | 0.815 | | *** | | | | 0.277 | | *** | | | | 0.323 | | *** | |
|  | | Detritusphere vs. Root rhizoplane | | 0.324 | | *** | | | | 0.515 | | *** | | | | 0.432 | | *** | | | | 0.425 | | *** | |
|  | | Detritusphere vs. Rhizosphere soil | | 0.367 | | *** | | | | 0.812 | | *** | | | | 0.312 | | *** | | | | 0.364 | | *** | |
|  | | Detritusphere vs. Root zone soil | | 0.384 | | *** | | | | 0.855 | | *** | | | | 0.319 | | *** | | | | 0.369 | | *** | |
|  | | Root rhizoplane vs. Rhizosphere soil | | 0.260 | | *** | | | | 0.525 | | *** | | | | 0.478 | | *** | | | | 0.580 | | *** | |
|  | | Root rhizoplane vs. Root zone soil | | 0.273 | | *** | | | | 0.574 | | *** | | | | 0.500 | | 0.015* | | | | 0.592 | | *** | |
|  | | Rhizosphere soil vs. Root zone soil | | 0.055 | | 0.004** | | | | 0.252 | | *** | | | | 0.252 | | *** | | | | 0.079 | | 0.023* | |
|  | | Phylloplane vs. Bulk soil | | 0.382 | | *** | | | | 0.752 | | *** | | | | 0.249 | | *** | | | | 0.303 | | *** | |
|  | | Detritusphere vs. Bulk soil | | 0.366 | | *** | | | | 0.787 | | *** | | | | 0.265 | | *** | | | | 0.296 | | *** | |
|  | | Root rhizoplane vs. Bulk soil | | 0.301 | | *** | | | | 0.578 | | *** | | | | 0.578 | | *** | | | | 0.605 | | *** | |
|  | | Rhizosphere soil vs. Bulk soil | | 0.224 | | *** | | | | 0.681 | | *** | | | | 0.315 | | *** | | | | 0.348 | | *** | |
|  | | Root zone soil vs. Bulk soil | | 0.225 | | *** | | | | 0.721 | | *** | | | | 0.305 | | *** | | | | 0.309 | | *** | |
| *CM* | | Phylloplane vs. Detritusphere | | 0.253 | | *** | | | | 0.401 | | *** | | | | 0.393 | | *** | | | | 0.492 | | *** | |
|  | | Phylloplane vs. Root rhizoplane | | 0.305 | | *** | | | | 0.436 | | *** | | | | 0.570 | | *** | | | | 0.640 | | *** | |
|  | | Phylloplane vs. Rhizosphere soil | | 0.277 | | *** | | | | 0.539 | | *** | | | | 0.417 | | *** | | | | 0.397 | | *** | |
|  | | Phylloplane vs. Root zone soil | | 0.269 | | *** | | | | 0.553 | | *** | | | | 0.328 | | *** | | | | 0.299 | | *** | |
|  | | Detritusphere vs. Root rhizoplane | | 0.378 | | *** | | | | 0.662 | | *** | | | | 0.521 | | *** | | | | 0.565 | | *** | |
|  | | Detritusphere vs. Rhizosphere soil | | 0.374 | | *** | | | | 0.852 | | *** | | | | 0.407 | | *** | | | | 0.480 | | *** | |
|  | | Detritusphere vs. Root zone soil | | 0.364 | | *** | | | | 0.862 | | *** | | | | 0.355 | | *** | | | | 0.444 | | *** | |
|  | | Root rhizoplane vs. Rhizosphere soil | | 0.270 | | *** | | | | 0.628 | | *** | | | | 0.562 | | *** | | | | 0.574 | | *** | |
|  | | Root rhizoplane vs. Root zone soil | | 0.284 | | *** | | | | 0.658 | | *** | | | | 0.345 | | 0.055 | | | | 0.536 | | *** | |
|  | | Rhizosphere soil vs. Root zone soil | | 0.055 | | 0.002** | | | | 0.179 | | *** | | | | 0.179 | | *** | | | | 0.098 | | 0.026* | |
|  | | Phylloplane vs. Bulk soil | | 0.308 | | *** | | | | 0.525 | | *** | | | | 0.282 | | *** | | | | 0.297 | | *** | |
|  | | Detritusphere vs. Bulk soil | | 0.382 | | *** | | | | 0.821 | | *** | | | | 0.390 | | *** | | | | 0.519 | | *** | |
|  | | Root rhizoplane vs. Bulk soil | | 0.346 | | *** | | | | 0.682 | | *** | | | | 0.552 | | *** | | | | 0.643 | | *** | |
|  | | Rhizosphere soil vs. Bulk soil | | 0.199 | | *** | | | | 0.642 | | *** | | | | 0.345 | | *** | | | | 0.398 | | *** | |
|  | | Root zone soil vs. Bulk soil | | 0.178 | | *** | | | | 0.607 | | *** | | | | 0.341 | | *** | | | | 0.240 | | *** | |
| *HM* | | Phylloplane vs. Detritusphere | | 0.217 | | *** | | | | 0.358 | | *** | | | | 0.279 | | *** | | | | 0.290 | | *** | |
|  | | Phylloplane vs. Root rhizoplane | | 0.298 | | *** | | | | 0.438 | | *** | | | | 0.198 | | *** | | | | 0.201 | | *** | |
|  | | Phylloplane vs. Rhizosphere soil | | 0.274 | | *** | | | | 0.521 | | *** | | | | 0.276 | | *** | | | | 0.315 | | *** | |
|  | | Phylloplane vs. Root zone soil | | 0.282 | | *** | | | | 0.550 | | *** | | | | 0.262 | | *** | | | | 0.282 | | *** | |
|  | | Detritusphere vs. Root rhizoplane | | 0.413 | | *** | | | | 0.682 | | *** | | | | 0.513 | | *** | | | | 0.586 | | *** | |
|  | | Detritusphere vs. Rhizosphere soil | | 0.361 | | *** | | | | 0.836 | | *** | | | | 0.670 | | *** | | | | 0.755 | | *** | |
|  | | Detritusphere vs. Root zone soil | | 0.376 | | *** | | | | 0.856 | | *** | | | | 0.680 | | *** | | | | 0.781 | | *** | |
|  | | Root rhizoplane vs. Rhizosphere soil | | 0.343 | | *** | | | | 0.701 | | *** | | | | 0.340 | | *** | | | | 0.418 | | *** | |
|  | | Root rhizoplane vs. Root zone soil | | 0.373 | | *** | | | | 0.741 | | *** | | | | 0.359 | | *** | | | | 0.426 | | *** | |
|  | | Rhizosphere soil vs. Root zone soil | | 0.072 | | *** | | | | 0.269 | | *** | | | | 0.147 | | *** | | | | 0.217 | | *** | |
|  | | Phylloplane vs. Bulk soil | | 0.273 | | *** | | | | 0.495 | | *** | | | | 0.214 | | *** | | | | 0.239 | | *** | |
|  | | Detritusphere vs. Bulk soil | | 0.361 | | *** | | | | 0.801 | | *** | | | | 0.627 | | *** | | | | 0.778 | | *** | |
|  | | Root rhizoplane vs. Bulk soil | | 0.398 | | *** | | | | 0.737 | | *** | | | | 0.349 | | *** | | | | 0.434 | | *** | |
|  | | Rhizosphere soil vs. Bulk soil | | 0.218 | | *** | | | | 0.676 | | *** | | | | 0.350 | | *** | | | | 0.475 | | *** | |
|  | | Root zone soil vs. Bulk soil | | 0.211 | | *** | | | | 0.673 | | *** | | | | 0.275 | | *** | | | | 0.314 | | *** | |
| *SP* | | Phylloplane vs. Detritusphere | | 0.290 | | *** | | | | 0.404 | | *** | | | | 0.232 | | *** | | | | 0.263 | | *** | |
|  | | Phylloplane vs. Root rhizoplane | | 0.266 | | *** | | | | 0.448 | | *** | | | | 0.356 | | *** | | | | 0.410 | | *** | |
|  | | Phylloplane vs. Rhizosphere soil | | 0.271 | | *** | | | | 0.563 | | *** | | | | 0.301 | | *** | | | | 0.327 | | *** | |
|  | | Phylloplane vs. Root zone soil | | 0.288 | | *** | | | | 0.600 | | *** | | | | 0.380 | | *** | | | | 0.420 | | *** | |
|  | | Detritusphere vs. Root rhizoplane | | 0.378 | | *** | | | | 0.624 | | *** | | | | 0.463 | | *** | | | | 0.521 | | *** | |
|  | | Detritusphere vs. Rhizosphere soil | | 0.370 | | *** | | | | 0.800 | | *** | | | | 0.335 | | *** | | | | 0.255 | | *** | |
|  | | Detritusphere vs. Root zone soil | | 0.396 | | *** | | | | 0.842 | | *** | | | | 0.424 | | *** | | | | 0.338 | | *** | |
|  | | Root rhizoplane vs. Rhizosphere soil | | 0.275 | | *** | | | | 0.594 | | *** | | | | 0.475 | | *** | | | | 0.520 | | *** | |
|  | | Root rhizoplane vs. Root zone soil | | 0.317 | | *** | | | | 0.676 | | *** | | | | 0.569 | | *** | | | | 0.626 | | *** | |
|  | | Rhizosphere soil vs. Root zone soil | | 0.121 | | *** | | | | 0.524 | | *** | | | | 0.044 | | 0.388 | | | | 0.040 | | 0.477 | |
|  | | Phylloplane vs. Bulk soil | | 0.276 | | *** | | | | 0.523 | | *** | | | | 0.292 | | *** | | | | 0.330 | | *** | |
|  | | Detritusphere vs. Bulk soil | | 0.394 | | *** | | | | 0.780 | | *** | | | | 0.309 | | *** | | | | 0.289 | | *** | |
|  | | Root rhizoplane vs. Bulk soil | | 0.344 | | *** | | | | 0.677 | | *** | | | | 0.445 | | *** | | | | 0.564 | | *** | |
|  | | Rhizosphere soil vs. Bulk soil | | 0.221 | | *** | | | | 0.700 | | *** | | | | 0.340 | | *** | | | | 0.376 | | *** | |
|  | | Root zone soil vs. Bulk soil | | 0.231 | | *** | | | | 0.725 | | *** | | | | 0.427 | | *** | | | | 0.502 | | *** | |

Note: *AO* *A. ordosica*; *CK* *C*. *korshinskii*; *HM* *H*. *mongolicum*; *SP* *S*. *psammophila*. *R*^2^ Adonis test statistic. Significant levels: **p* < 0.05, ** *p* < 0.01, *** *p* < 0.001.

**Table S5** Microbial co-occurrence network characteristics in different microhabitats.

| Domains | Habitats | Node | Edge | AD | Modularity | ACC | APD | Hub node |
| --- | --- | --- | --- | --- | --- | --- | --- | --- |
| **Bacteria**  **Fungi** | Bulk soil  Root zone soil  Rhizosphere soil | 963  1167  1138 | 20723  2681  2624 | 43.038  4.595  4.612 | 0.911  0.744  0.740 | 0.997  0.303  0.270 | 1.000  5.740  5.767 | 697  0  0 |
|  | Root rhizoplane | 600 | 1596 | 5.320 | 0.751 | 0.364 | 5.462 | 0 |
|  | Detritusphere | 537 | 846 | 3.151 | 0.803 | 0.258 | 7.082 | 0 |
|  | Phylloplane  Bulk soil  Root zone soil  Rhizosphere soil  Root rhizoplane  Detritusphere  Phylloplane | 958  510  434  442  417  294  547 | 5687  7761  2434  1729  17214  2082  4841 | 11.873  30.435  5.608  3.912  41.281  14.163  17.700 | 0.768  0.849  0.768  0.721  0.328  1.043  0.760 | 0.477  0.991  0.381  0.347  0.525  0.489  0.588 | 5.850  1.016  8.763  5.745  3.070  4.056  5.025 | 0  341  0  0  157  51  0 |

AD: average degree; ACC: average clustering coefficient; APD: average path distance

**Table S6** Percentage of nodes from different bacterial and fungal phyla in the co-occurrence network.

| Domain | Phylum | Bulk soil | Root zone soil | Rhizosphere soil | Root rhizoplane | Detritusphere | Phylloplane |
| --- | --- | --- | --- | --- | --- | --- | --- |
| **Bacteria** | Proteobacteria  Actinobacteria  Planctomycetes  Bacteroidetes  Acidobacteria  Chloroflexi  Patescibacteria  Gemmatimonadetes  Others | 20.25%  14.33%  15.06%  6.44%  6.85%  9.55%  6.44%  4.98%  16.10% | 19.71%  14.05%  15.85%  8.31%  6.94%  11.74%  6.34%  5.40%  11.66% | 21.35%  14.29%  15.29%  11.34%  7.21%  11.42%  3.95%  4.48%  10.67% | 32.83%  12.83%  9.83%  17.17%  4.50%  6.17%  4.00%  2.00%  10.67% | 30.91%  9.68%  7.82%  28.68%  3.91%  2.98%  1.49%  2.23%  12.30% | 29.12%  10.13%  9.50%  18.58%  6.37%  3.97%  2.51%  2.09%  17.73% |
| **Fungi** | Ascomycota  Basidiomycota  Zygomycota  Chytridiomycota  Glomeromycota  Unidentified  Others | 57.25%  17.65%  2.35%  2.75%  1.57%  18.04%  0.039% | 61.29%  16.13%  2.30%  2.30%  3.46%  14.29%  0.23% | 62.44%  16.74%  2.94%  1.58%  1.58%  14.48%  0.24% | 56.83%  30.46%  1.68%  0.72%  0.96%  8.63%  0.72% | 66.33%  18.71%  0%  0.34%  0%  13.95%  0.67% | 57.59%  22.49%  1.10%  0.73%  0.73%  15.36%  2.00% |
